# Locating causal hubs of memory consolidation in spontaneous brain network

**DOI:** 10.1101/2022.05.08.490699

**Authors:** Zengmin Li, Dilsher Athwal, Hsu-Lei Lee, Pankaj Sah, Patricio Opazo, Kai-Hsiang Chuang

## Abstract

Memory consolidation after learning involves spontaneous, brain-wide network reorganization during rest and sleep, but how this is achieved is still poorly understood. Current theory suggests that the hippocampus is pivotal for reshaping the connectivity. Here we identify that a different set of spontaneous networks and their hubs are instrumental in consolidating memory during post-learning rest. We found that two types of spatial memory training invoke distinct functional connections, but a network of the sensory cortex and subcortical areas is common for both tasks. Furthermore, learning increased brain-wide network integration, with the prefrontal, striatal and thalamic areas being influential for this network-level reconfiguration. Chemogenetic suppression of each hub identified after learning resulted in retrograde amnesia, confirming the behavioral significance. These results demonstrate the causal and functional roles of resting-state network hubs in memory consolidation and suggest a distributed network beyond the hippocampus subserving this process.

## INTRODUCTION

The formation of enduring memory in the brain is a distributed and dynamic process, the mechanism of which is not fully understood. Current theory of systems memory consolidation suggests that the hippocampus mediates the encoding of information from segregated sensory, motor, or motivation brain networks that are engaged during learning, gradually reshaping their connectivity to form long-term memory ^1,2^. This is facilitated by the reactivation of learning-associated neuronal populations (replay) and the coordinated interaction of the hippocampal-neocortical network during post-encoding periods of quiet wakefulness (resting state) and sleep ^3–6^. Apart from the hippocampus, where, when and how other regions are involved in facilitating this system-wide reconfiguration are still unclear. Whole-brain functional imaging during task performance has revealed broad engagement of not only neocortical but also subcortical areas when encoding or recalling memory in humans ^7^ and rodents ^8,9^. However, the regions involved in consolidating memory in the ill-defined, “offline” period are difficult to pinpoint without aligning them to activities associated with replay ^10^.

A major advance over the last decade has been the identification of the brain-wide network involved in spontaneous activity during task-free conditions ^11^. Functional connectivity (FC), measured as the correlation between regional activities at resting state, forms large-scale networks of functionally associated areas that indicate an intrinsic organization of the brain ^12,13^. The disruption of these resting-state networks (RSNs) in association with cognitive impairment in aging and disease provides evidence for their involvement in the etiology and progression of these conditions ^14–16^. Advances in network neuroscience have further revealed that the topology of RSNs alters with performance or symptom, suggesting the behavioral relevance of their organization and dynamics ^17,18^. Importantly, learning can induce ongoing remodeling of the RSNs over time in humans ^19,20^ and rodents ^21,22^. Increased association between hippocampus-neocortical FC and performance over repeated training ^23,24^, reconfiguration after sleep ^25,26^, and reactivation of the learning-related activity pattern ^27,28^ suggested that post-encoding RSNs reflect systems memory consolidation.

However, a fundamental question that remains is whether the spontaneous network changes are causative of the behavior- or disease-associated states with which they correlate ^29^. Due to the observational nature of the experimental designs, unconstrained imaging environments, and correlation-based FC measures, it remains possible that the RSN changes are epiphenomena that are driven by alternative neural, physiological or pathological factors ^30–34^. Furthermore, if they are causative, what activity and which area drive such large-scale network remodeling is unresolved. Analytical methods can allow the inference of causality from the RSNs (for review, see ^35^). Nonetheless, they only estimate inter-dependency between regional activities within a network, instead of the causality to behavior. Critically, whether a network or its hub is causally required for cognition (e.g., episodic memory), such that its dysfunction leads to a disability (e.g., amnesia) whereas its facilitation improves performance, has not been directly tested experimentally with prospective interventions.

In this study, we reveal the brain networks that are instrumental in consolidating memory during post-encoding rest by identifying and functionally manipulating RSN hubs. We examined two hypotheses for defining causal hubs of behavior, one based on a common network and the other on network integration. Certain elements of the hippocampal-neocortical network, particularly between the hippocampal formation (HPF, including the hippocampus, subiculum and entorhinal cortex), retrosplenial cortex and medial prefrontal cortex (mPFC), have been identified in different spatial or contextual learning paradigms ^36–40^. The storage of various forms of the spatial memory trace (engram) in these areas ^38^ indicates that a common, task-invariant network may be involved in the systems consolidation ^1,41^. Yet, the full extent of this common network is still unclear. In addition, consolidation incorporates new information from functionally segregated areas, such that this integration could be manifested at the network level. Indeed, topological features of network integration, such as global efficiency ^42^, or segregation, such as modularity ^43^, have been shown to correlate with performance during or after learning. Altered network integration is also found after cognitively demanding tasks ^44,45^ or overnight consolidation ^25^. These findings indicate that network integration is an essential feature in learning and memory. Thus, influential hubs for network integration (“integrator” hubs) may have a causal role in memory formation.

To test whether common or integrator hubs are causally involved in memory consolidation, we trained mice in two versions (early and late retrieval) of active place avoidance (APA), a spatial memory task, and subsequently acquired resting-state functional magnetic resonance imaging (rsfMRI) data to characterize behavior-induced RSN changes (Fig. 1a). We developed methods for detecting common or integrator hubs from post-encoding RSNs. We then validated the behavioral impact of the identified hubs by silencing each hub individually during the consolidation period using inhibitory Designer Receptors Exclusively Activated by Designer Drugs (DREADDs) ^46^.

**Figure 1.**
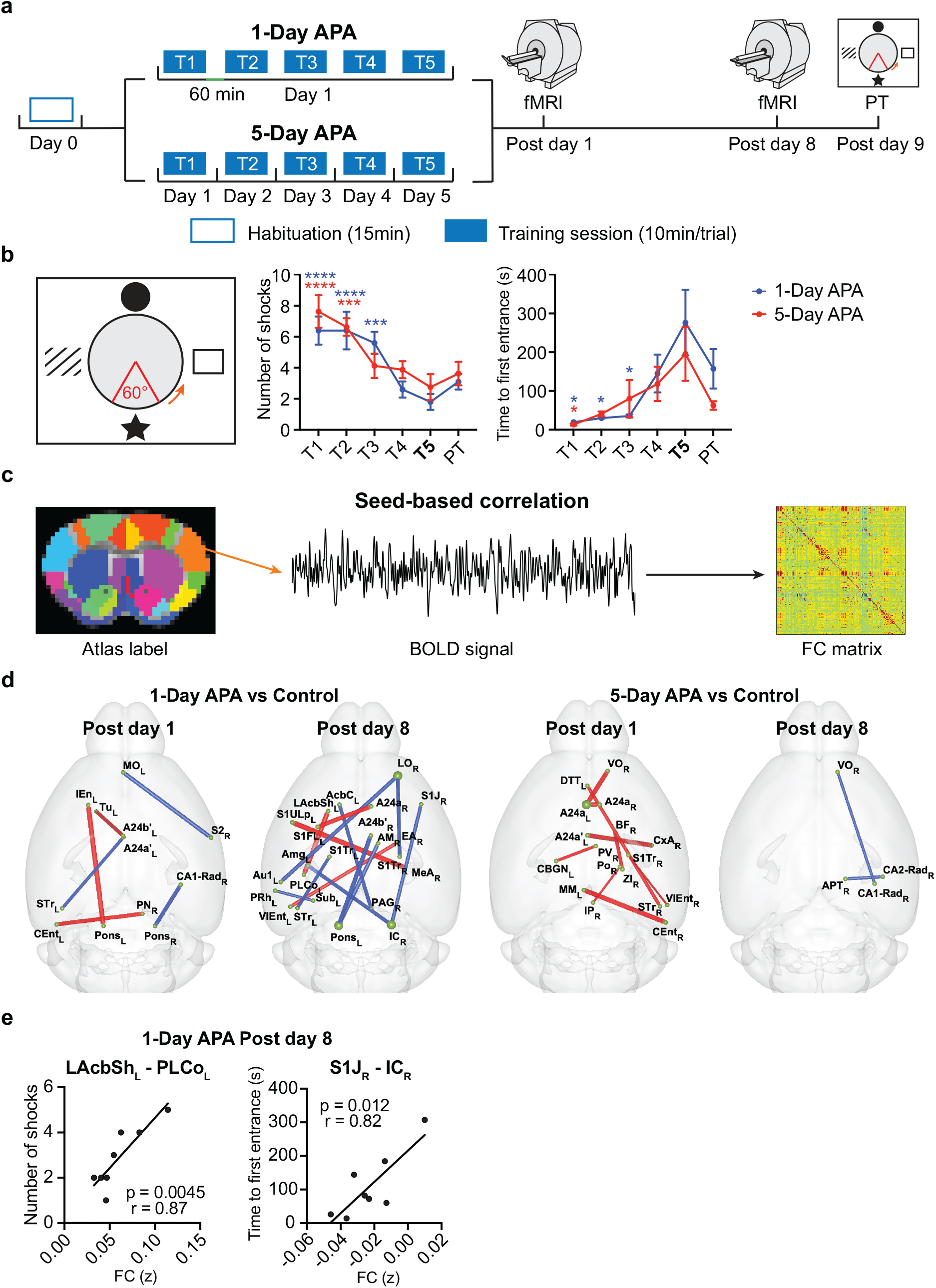
RSN changes after spatial learning in mice. (**a**) The schematic diagram of the APA-rsfMRI experiment. (**b**) The left diagram shows the setup of the APA task. Four distinct pictures were hung on the surrounding walls as visual cues. The orange arrow indicates the direction of rotation. The 60° sector in red shows the location of the invisible “shock zone”. The two plots on the right show the progressive decrease in the number of shocks over the trials (two-way ANOVA, F_5, 80_ = 14.22, p < 0.0001), which is comparable between the 1-Day (N = 10) and 5-Day APA (N = 8) groups (F_1, 16_ = 0.55, p = 0.47 for groups). Similar trends can be seen in the time to first entrance of the shock zone (two-way ANOVA, F_5, 80_ = 6.63, p < 0.0001 for training trials, F_1, 16_ = 1.27, p = 0.28 for groups). Comparisons were performed between the last training trial (T5) and other training trials (T1 to T4) or probe test (PT). Data are represented as mean ± SEM. * p < 0.05; *** p < 0.001; **** p<0.0001. (**c**) The seed-based correlation analysis used to create the FC matrix of each animal. (**d**) Changed functional connections in the 1-Day and 5-Day APA, compared to their corresponding controls, on post-training day 1 and post-training day 8 (two-sample t-test, p < 0.05, FDR corrected; see Supplementary Table S3 for N of each group). The node size indicates the degree of the node. The red connections represent APA > control while the blue connections represent APA < control. The line thickness indicates the t value. (**e**) Two functional connections from the 1-Day APA post-training day 8 correlated with the memory retention probe test (N = 8).

## RESULTS

### Similar behavior leads to distinct post-encoding RSNs

It has been shown that the HPF, mPFC and retrosplenial cortex are typically involved in the consolidation of spatial or contextual memory. To investigate whether similar spatial learning invokes a common network in post-encoding rest, we conducted two APA tasks with the same training trials but different inter-trial intervals: one hour (1-Day APA, as the learning was completed in one day) versus one day (5-Day APA; Fig. 1a). APA allows spatial navigation and memory to be assessed in mice with less stress than that associated with the water maze by training them to avoid a shock zone based on spatial cues over repeated trials (Fig. 1b) ^47,48^. After learning, two sessions of rsfMRI were performed on post-training days 1 and 8 to examine the plasticity of the RSNs. One day after the second rsfMRI scan, a probe test was performed to assess memory retention. The number of shocks (N_shock_) that the animals received and the time to first entrance into the shock zone (T_enter_) were used to measure their behavioral performance. N_shock_ gradually decreased and T_enter_ increased during learning; this, together with the similar values obtained during the probe tests (Fig. 1b; Supplementary Data), demonstrated that the mice could remember both APA tasks equally well after 9 days, demonstrating the formation of long-lasting memory.

We distinguished post-encoding RSNs by comparing the FC between 230 highly parcellated brain regions in a brain template (Fig. 1c; Supplementary Table S1) of the APA groups versus their controls. In the control group, animals were exposed to the APA training procedures without any foot shock being delivered as we found that random shock elicited a strong stress response. Despite comparable behaviors during learning and retrieval, we found distinct post-encoding RSNs. On post-training day 1 (Fig. 1d; two-sample t-tests, p < 0.05, false discovery rate [FDR]-corrected), the 1-Day APA increased sparse FC in the left hemisphere between the entorhinal cortex and motor control area (pontine nucleus), and between reward-processing areas (olfactory tubercle and dorsal anterior cingulate cortex [A24a], part of the mPFC), but decreased the FC between the somatosensory and prefrontal cortices and between the HPF and pons in the right hemisphere. In contrast, the 5-Day APA increased the FC mostly in the right hemisphere, including the HPF, prefrontal cortex and sensory areas. This highly lateralized FC (12 out of 17 connections) is consistent with studies reporting that the right hemisphere is predominant in memory processing ^49–51^. One week after APA training, the network reorganized. In the 1-Day APA group, even more inter-hemispheric FC was found, with increased FC being observed among the somatosensory cortex, areas for emotional and motor responses (e.g., lateral accumbens shell [LAcbSh]) and ventral entorhinal cortex, whereas the FC between the lateral orbital cortex (LO, part of the prefrontal cortex), somatosensory cortex, thalamus, and pons decreased. In the 5-Day APA group, only sparse FC between the HPF, prefrontal cortex and thalamus in the right hemisphere was found. These results indicated that post-encoding RSNs are task and time dependent and involve distant neocortical and subcortical areas, similar to the findings of previous studies using the Morris water maze ^21,22^. The overall connectivity with the HPF and mPFC is consistent with their critical roles in memory consolidation, although the specific subregions involved differed between tasks. A common network between the two APA tasks could be obscured with such detailed parcellation. Alternatively, this could be due to the much stricter false positive rate when calculating the overlap between two FDR-corrected connectivity matrices.

It is generally expected that behaviorally relevant connections predict performance. Many studies have found an association between memory performance and FC during encoding or retrieval ^52–54^, yet little is known about the relationship with post-encoding FC. To test whether post-encoding FC can predict memory retention, we calculated Pearson’s correlation between FC strength and N_shock_ or T_enter_ in the probe trial (Fig. 1e). Only two connections on post-traning day 8 in the 1-Day APA group correlated with behavior: between the left posterolateral cortical amygdala (PLCo) and the LAcbSh (r = 0.87, p = 0.0045) and between the right primary somatosensory cortex jaw region and the inferior colliculus (r = 0.82, p = 0.012). Although several connections were enhanced in the 5-Day APA, no correlation with behavior was found. This indicates that the most significant FC may not be influential on behavior.

### Locating common network hubs that predict behavior

We predicted that RSNs commonly induced by both kinds of APA tasks are influential for memory consolidation. To discover common RSNs induced by both tasks, the threshold for the two-sample t-tests was lowered to an uncorrected p<0.05 (Fig. 2a). Despite being similar in their task designs, only a small fraction of the FC was common in both APA tasks, with 3.56% (post-training day 1) and 2.85% (post-training day 8) of the connections overlapping. To identify the causal hub, we selected the common FC that was predictive of behavioral performance in the probe test. Using a permutation test, imposing behavioral correlation with N_shock_ (threshold at p<0.05) resulted in an equivalent family-wise error of p < 0.05 for common connections on both post-training day 1 and 8 (Fig. 2b,c). When using T_enter_ as a behavioral index, the family-wise false positive rate of common connections was p = 0.019 on posttraining day 1 but was not significant for post-training day 8 (Supplementary Fig. S1). Here, we chose to use N_shock_ as the primary behavioral index.

**Figure 2.**
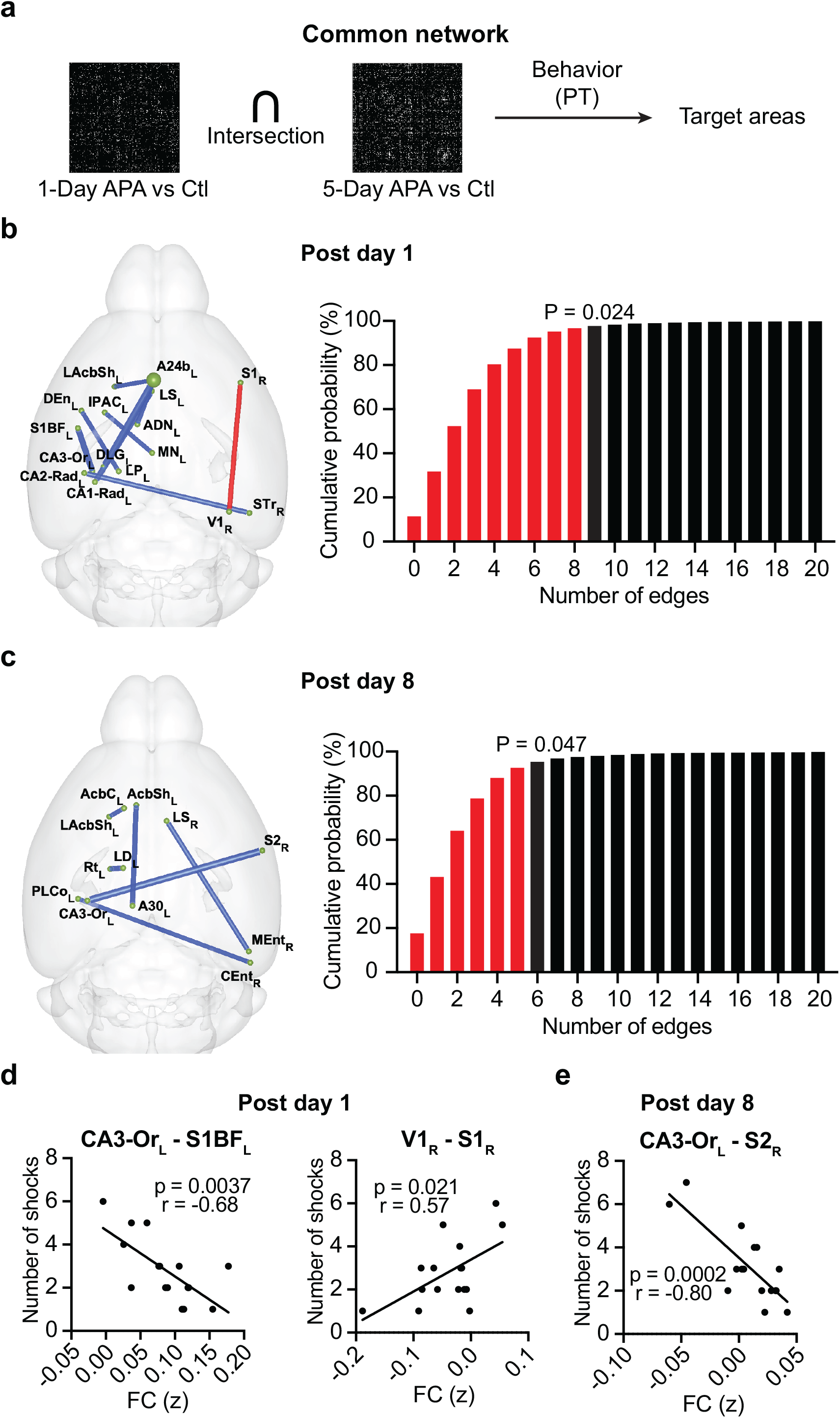
Identification of common networks. (**a**) Procedures for identifying target hubs based on common networks that predict (correlate) behaviors in the probe test (PT). The difference FC matrix between the APA and control (Ctl) groups was calculated by two-sample t-test. The FC on (**b**) post-training day 1 (N = 17) and (**c**) post-training day 8 (N = 16) that correlated with N_shock_ (p<0.05, two-tailed) within the common networks (p<0.05, uncorrected) of the 1-Day and 5-Day APA. The node size indicates the degree of the node. The red lines show the functional connections that positively correlate with N_shock_ while the blue lines show those that negatively correlate with N_shock_. The line width indicates the absolute r value. The cumulative distribution of the 5000 permutation tests is shown on the right. The red bars represent the cumulative probability before reaching the real number of edges (9 connections on post-training day 1 and 5 connections on posttraining day 8). Behavioral correlation of the functional connections with the target hubs on (**d**) post-training day 1 (N = 17) and (**e**) post-training day 8 (N = 16).

We found common networks composed of the hippocampus, mPFC and thalamus in the left hemisphere, and the connection between the primary somatosensory and primary visual (V1) cortex in the right hemisphere on post-training day 1 (Fig. 2b, Supplementary Table S2). Excluding the HPF and mPFC, which are known to be engaged in memory consolidation, and subcortical areas, the FC of the left primary somatosensory cortex barrel field (S1BF_L_) had the highest behavioral correlation (r = −0.68), followed by right V1 (r = 0.57; Fig. 2d). On posttraining day 8 (Fig. 2c, Supplementary Table S2), the FC with the mPFC was gone. Instead, we observed FC with the reticular nucleus (Rt), which drives the neural oscillations important for memory consolidation during sleep ^5^, and the retrosplenial cortex. The engagement of mPFC on day 1 but silencing one week later is consistent with the temporal dynamic of the engram in this area ^38^. Excluding the entorhinal and retrosplenial cortices, the right secondary somatosensory cortex (S2_R_) had the highest behavioral correlation (r = −0.80, Fig. 2e). Overall, we found expanded common networks beyond the HPF, mPFC and retrosplenial cortex were engaged at different time after learning. Here, we chose the S1BF_L_, V1_R_ and S2_R_ as the targets for validation.

### Learning alters network integration

Based on the importance of network integration in learning and memory ^25,42,44,45,54^, we predicted that post-encoding RSNs are more integrated after spatial learning. To investigate this, we applied graph theory analysis, which simplifies the brain network as nodes (brain regions) and edges (FC strengths), to the connectivity matrices after two-sample t-tests. The global efficiency, modularity, and size of the giant component (the largest cluster of interconnected nodes which represents network integration) were used to evaluate the network integration and segregation. With increased t-score threshold from 2.0 to 3.8 (uncorrected), networks became more fragmented, resulting in a reduced giant component and global efficiency but increased modularity (Fig. 3a-c, and Supplementary Fig. S2). To test the overall difference, the area under the curve (AUC) was calculated ^55^ and compared to the distributions of 5000 random networks. Based on the null distribution, both APA training, except the 5-Day APA on post-training day 8, significantly increased the size of the giant component (Fig. 3a, and Supplementary Fig. S2a). The global efficiency was altered on post-training day 8 but not day 1 (Fig. 3b, and Supplementary Fig. S2b). Interestingly, the modularity was significantly increased in both APA tasks at both time points (Fig. 3c, and Supplementary Fig. S2c). This is consistent with a recent study which reported that repeated training that automates a cognitively demanding task can increase integration and segregation of post-encoding RSNs ^45^. With features of both network integration and segregation being increased, we examined the community properties that may cause such features. We found that a rapid increase in the proportion of connections within components but a plateau in the number of components led to the increased modularity (Supplementary Fig. S3). Together these results indicate that APA learning increases network integration and segregation by forming larger and more network components while also strengthening the connectivity within components.

**Figure 3.**
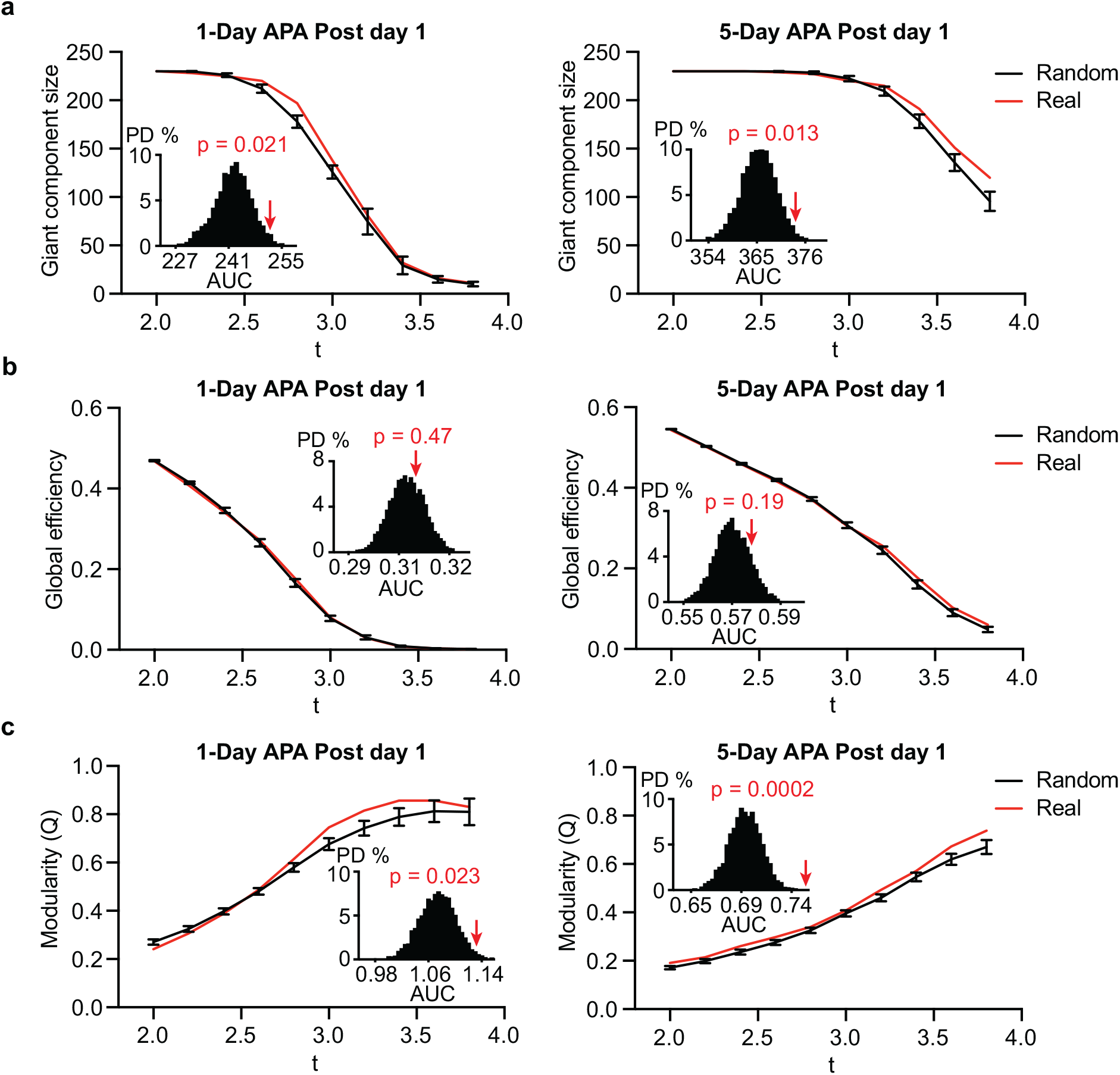
Graph characteristics of post-encoding RSNs. Trends of (**a**) giant component size, (**b**) global efficiency and (**c**) modularity of post-encoding RSNs for 1-Day (left) and 5-Day (right) APA on post-training day 1 (red) compared to random networks (black), thresholded at 2 ≤ t ≤ 3.8. The black lines show the mean ± SD of 5000 random networks generated based on the real network. The embedded bar graphs show the probability distribution of the area under curve (AUC) of the random networks. The red arrows indicate the AUC for the real network and the corresponding p value. PD: probability distribution.

### Optimal method for distinguishing integrator hubs

As network integration is a feature of post-encoding RSNs, pinpointing the integrator hubs would allow us to test whether this is causally required in memory consolidation. We predicted that when an integrator hub is removed (inhibited), the network integration would be greatly impeded, leading to the breakdown of the giant component. The best method for hub identification would be one which can reduce the size of the giant component with the removal of the fewest number of nodes. Centrality, which describes the importance of network communication and integration, is typically regarded as reflecting network integration. However, simulation showed that centrality is a poor measure of causal inference ^56^.

To determine the best method, we compared four centrality measures (degree centrality, closeness centrality, betweenness centrality, eigenvector centrality) and collective influence (CI), which searches for nodes that can quickly break down large networks in an optimal percolation model ^57,58^. We calculated the reduction in the giant component in the post-encoding RSNs by removing high ranking/centrality nodes one by one. Fig. 4b shows an example in which the normalized giant component size quickly dropped by removing nodes detected by the CI, followed by those identified by the degree centrality and betweenness centrality, whereas closeness and eigenvector centrality had a slower effect. Similar trends were observed in networks from both APA groups at both days 1 and 8 (Supplementary Fig. S4). Comparing the AUC of the giant component changes, the CI analysis resulted in the quickest collapse of the giant component (Fig. 4c, Supplementary Fig. S5). This indicates that CI is a better method for identifying integrator hubs.

**Figure 4.**
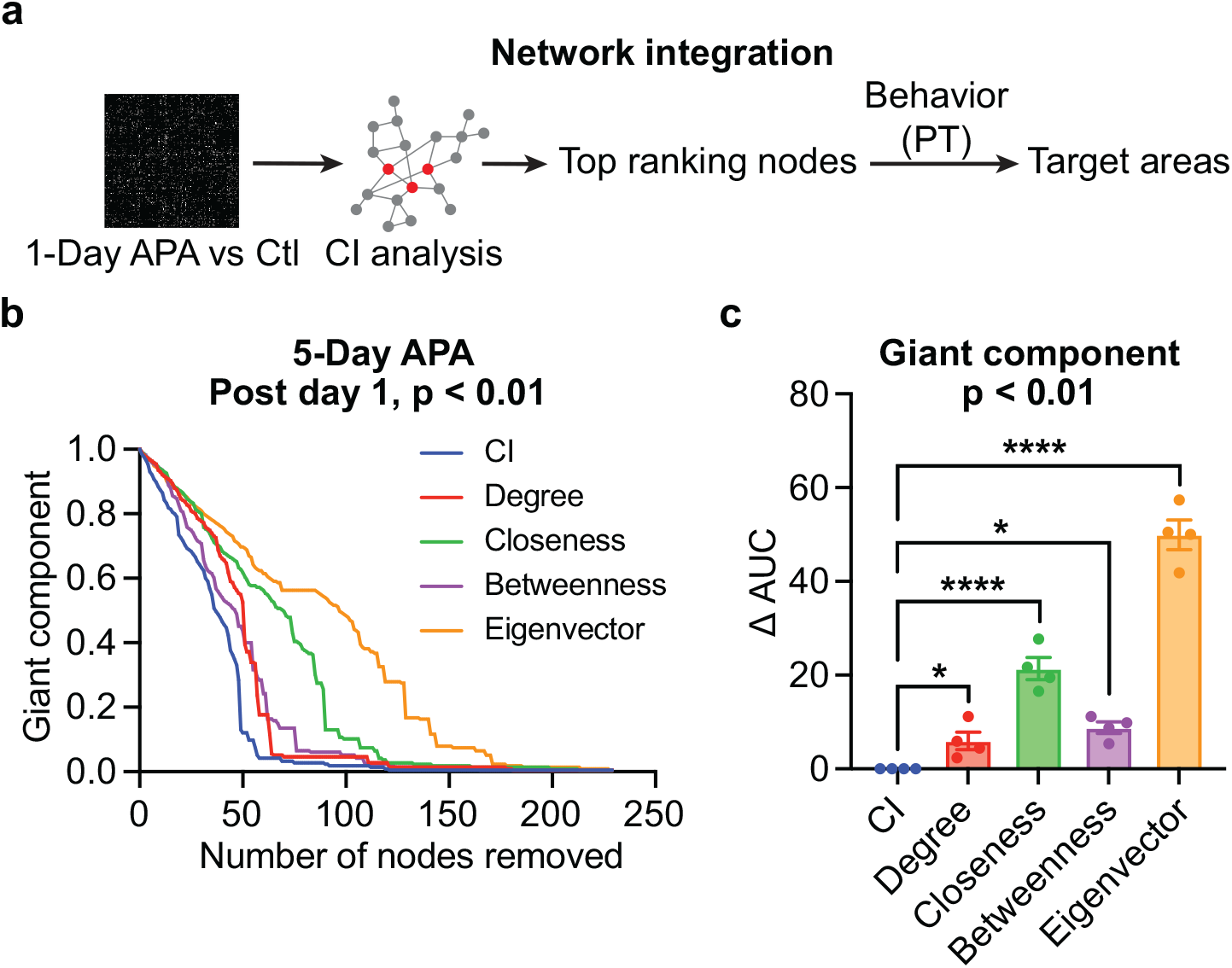
Comparison of hub selection methods for network integration. (**a**) Procedures for identifying target hubs based on network integration. The difference FC matrix between the APA and control (Ctl) groups was analyzed by CI to select top ranking nodes. Nodes with FC that predicts behaviors in the probe test (PT) were selected. (**b**) The relative giant component size decreases by removing high-ranking CI or centrality hubs one by one from the difference FC matrix between the 5-Day APA and its control on post-training day 1 (p<0.01, uncorrected). Five hub selection methods, including CI (blue), degree centrality (red), closeness centrality (green), betweenness centrality (purple), and eigenvector centrality (orange), were compared. The quicker the giant component shrinks, the more effective the original network breaks into smaller networks (reduced network integration). (**c**) Comparison of the area under the curve (AUC) of the giant component reduction curves. The data points include the difference FC matrices comparing APA and control groups for the 1-Day APA and 5-Day APA datasets on both post-training day 1 and day 8. Each data point was normalized by the corresponding AUC of CI. One-way ANOVA, F_4, 12_ = 122.3, p < 0.0001. Data are represented as mean ± SEM. * p < 0.05; **** p < 0.0001.

### Identification of integrator hubs that predict behavior

We next applied CI analysis to post-encoding RSNs of the 1-Day APA to identify nodes with FC that predicts memory retention (Fig. 4a). As shown in Fig. 1, the use of the FDR-corrected threshold makes the network very sparse without a giant component, thereby excluding the use of CI analysis for hub identification ^57,59^. Here, we used multiple uncorrected thresholds, p < 0.05, 0.01 and 0.005, to determine the averaged ranking of a node in breaking down the RNSs after 1-Day APA training. On post-training day 1, the 10 top-ranking nodes were mostly subcortical, including regions in the basal ganglia, midbrain and brainstem, with cortical areas in the HPF (subiculum and entorhinal cortex) and S2 being ranked lower (Table 1). On post-training day 8, more cortical areas (parietal association, sensory and prefrontal cortices) rose to the top ranking compared to subcortical areas. Considering the behavioral correlation of the FC with these CI nodes, only a few nodes had FC that was predictive of memory retention (Table 2). Our results showed that the FC of the right caudate putamen had the highest correlation with N_shock_ (r = −0.79, p = 0.0063), and the left LAchSh had the highest correlation (r = 0.95, p = 3.7×10^-5^) with T_enter_ on post-training day 1. On post-training day 8, nodes with high behavioral correlation include the left ventromedial thalamic nucleus (VM_L_, r = 0.91, p = 0.0015), left primary somatosensory cortex forelimb region (S1FL_L_; r = 0.89, p = 0.0033), right LO (r = 0.87, p = 0.0045) and right primary somatosensory cortex trunk region (S1TrR; r = −0.84, p = 0.0095). We therefore chose the LAchShL, LO_R_, VM_L_ and S1TrR as the top, middle and low ranking hubs for validation.

**Table 1.**
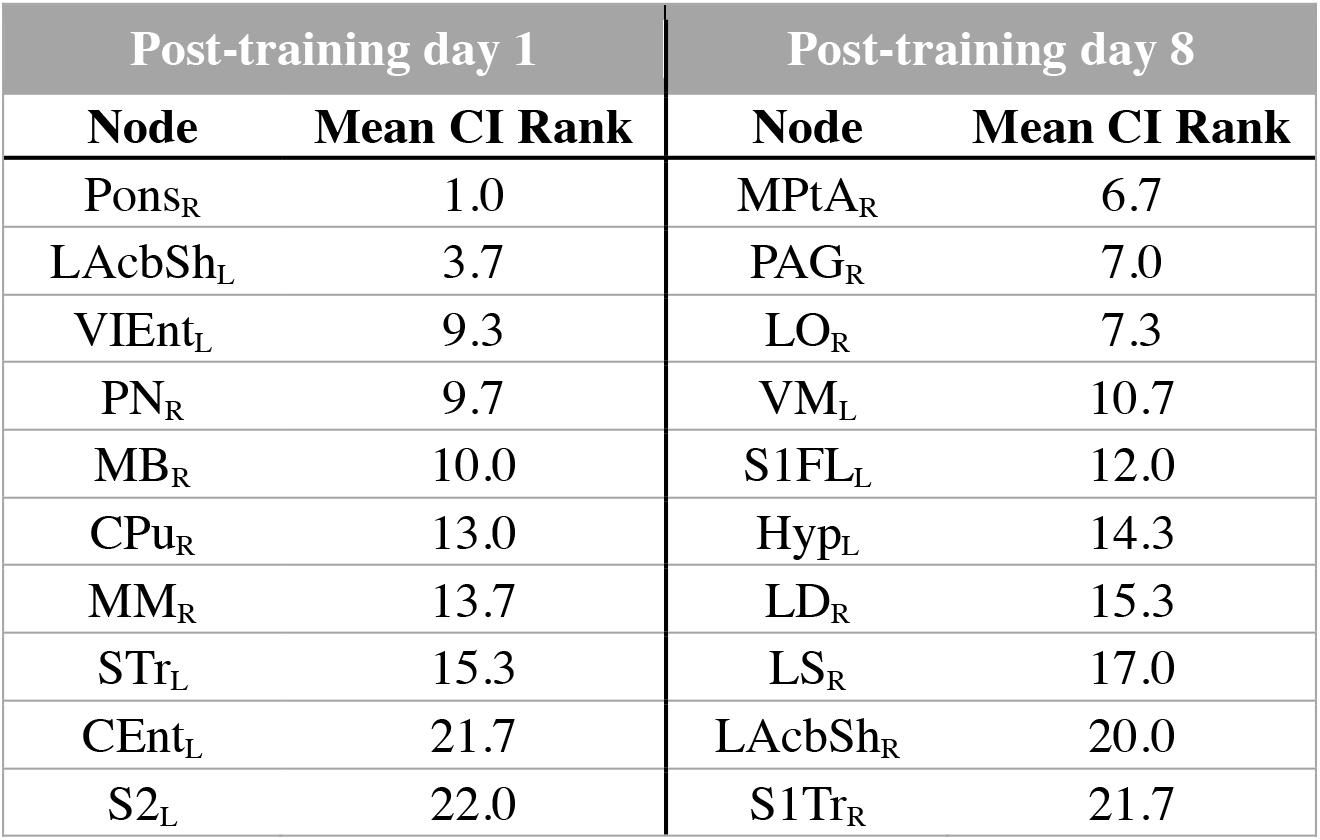
Top 10 nodes for network integration identified by CI analysis. This table shows the top 10 ranking nodes according to the mean CI rank under network threshold of p < 0.05, p < 0.01 and p < 0.005 when comparing the 1-Day APA and control. r: right hemisphere; l: left hemisphere. See Supplementary Table S1 for the abbreviations of brain regions.

**Table 2.**
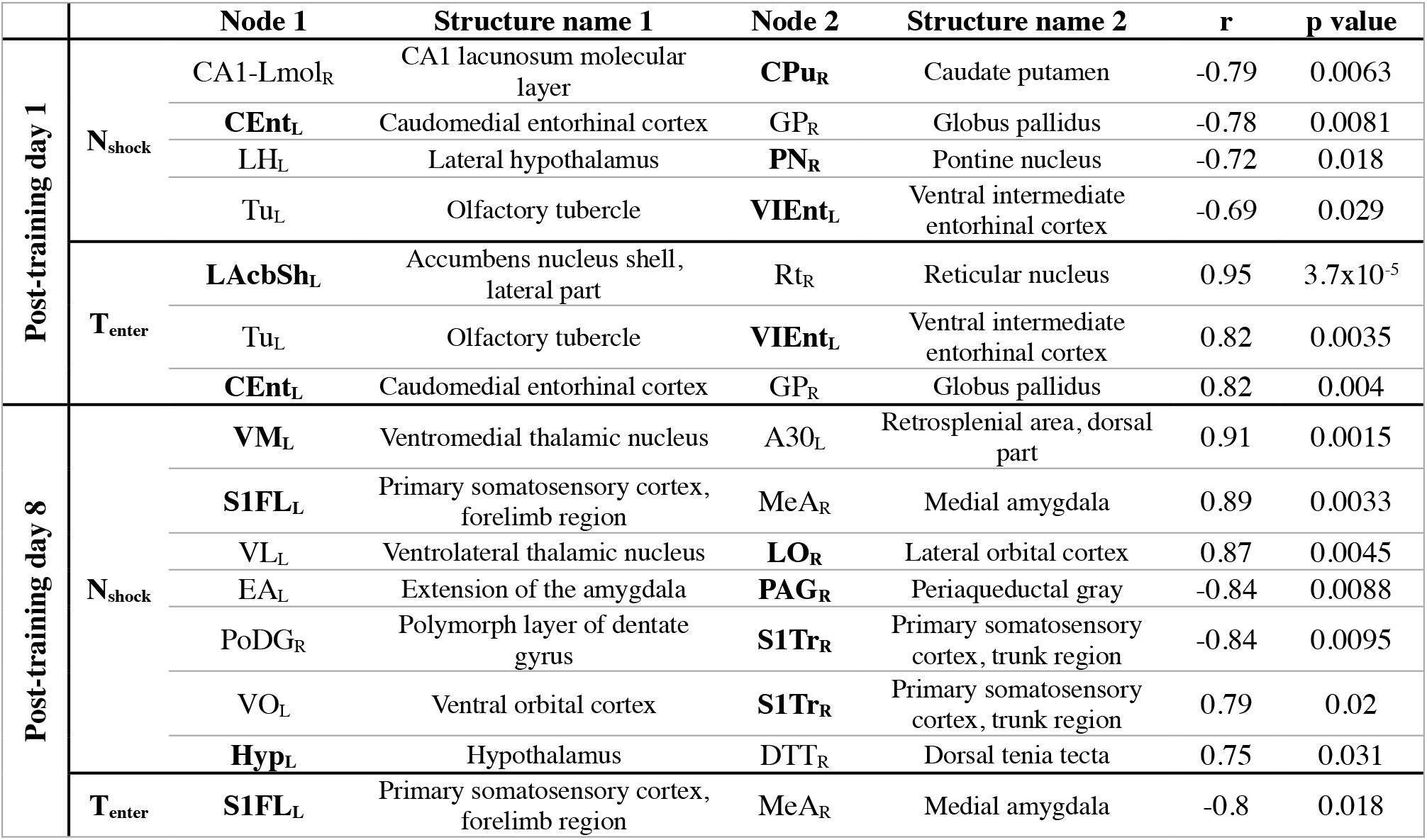
Behavior-correlated functional connections containing the top 10 CI nodes. Each row shows the names of the nodes, the behavioral correlation r and p values. Brain regions shown in **bold** are the top 10 ranking nodes in the CI analysis (shown in Table 1). r: right hemisphere; l: left hemisphere.

### Validation of causal hubs by DREADDs inhibition

To verify the causal role of selected hubs in memory consolidation, we injected AAV2/1-pSyn-hM4D(Gi)-T2A-mScarlet to transfect inhibitory DREADDs in all neurons in each area individually (Fig. 5a). One month after the surgery, animals went through the 1-Day APA training. Immediately after finishing the 5 training trials, animals were administered clozapine N-oxide (CNO) by intraperitoneal injection, followed by drinking water containing CNO to maintain inhibition of the targeted hubs for 7 days until one day before the probe test, to allow clearance of the CNO ^60,61^. In addition to a naïve group to control for the effect of CNO, we chose one cortical area, the right frontal association cortex (FrAR), and one subcortical area, the right ventral posteriomedial thalamus (VPM_R_), which did not change FC in our analyses, as negative controls. Fig. 5 illustrates good viral expression in the targeted areas in both the experimental and control groups, although we did notice that there was some viral expression in nearby brain areas, such as S2_R_ and LAcbShL.

**Figure 5.**
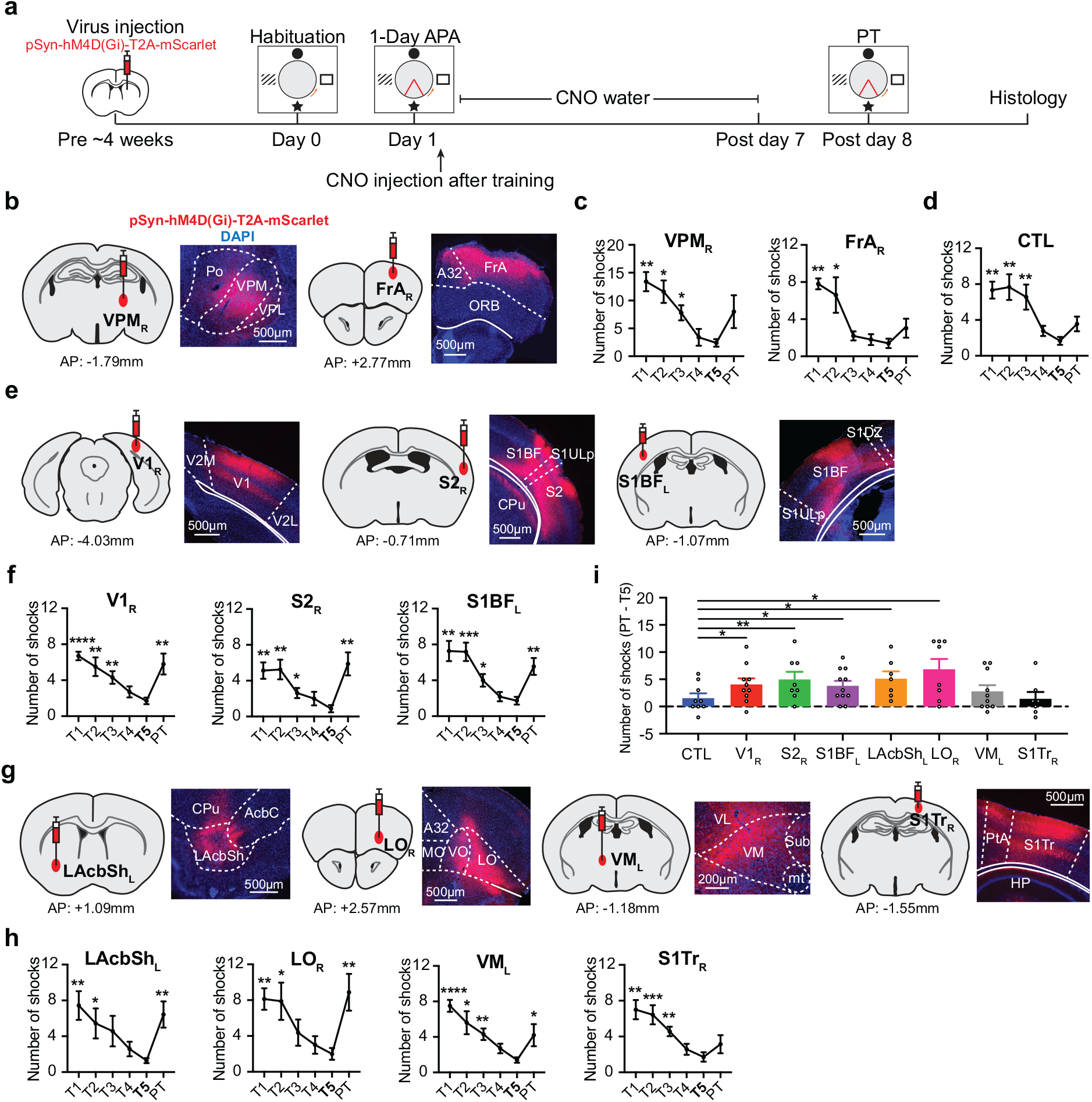
DREADDs expression and behavioral effects of hub inhibition. (**a**) Schematic diagram of target validation using DREADDs inhibition. Representative DREADDs expression (red) in the target areas selected from (**b**) the negative control, (**e**) common networks, and (**g**) network integration. In each subgraph, the left image shows the injection location of AAV-pSyn-hM4D(Gi)-T2A-mScarlet, and the right image shows the fluorescence imaging together with the DAPI staining (blue). (**c**) The number of shocks during the 1-Day APA learning trials (T1 to T5) show a progressive decrease and insignificant change during the probe test (PT) after inhibition of the negative controls, VPM_R_ (one-way ANOVA F_5, 20_ = 10.80, p < 0.0001; N = 5) and FrAR (F_5, 20_ = 10.65, p < 0.0001; N = 5). The post hoc comparison was conducted with respect to T5. (**d**) The behavior was comparable to that of the CNO-control (CTL) group (F_5, 40_ = 7.41, p < 0.0001; N = 9). (**f**) Similar trends during learning can be seen with DREADDs in the common hubs V1_R_ (F_5, 45_ = 6.95, p < 0.0001; N = 10), S2_R_ (F_5, 35_ = 6.02, p = 0.0004; N = 8) and S1BF_L_ (F_5, 50_ = 10.67, p < 0.0001; N = 11), or in the (**h**) integrator hubs LAchShL (F_5, 30_ = 5.31, p = 0.0031; N = 7), LO_R_ (F_5, 35_ = 6.13, p = 0.0004; N = 8), VM_L_ (F_5, 45_ = 7.16, p < 0.0001; N = 10) and S1TrR (F_5, 30_ = 6.40, p = 0.0004; N = 7). After hub inhibition during the 1-week consolidation period, impaired memory recall was seen in all the common and integrator hubs, except S1TrR. (**i**) Comparison of the change in the number of shocks (PT – T5) between the experimental groups and the CTL (two-sample t-test, one tail) shows an increased number of shocks after inhibition of S1BF_L_ (Cohen’s d = 0.76, p = 0.046), V1_R_ (Cohen’s d = 0.78, p = 0.045), S2_R_ (Cohen’s d = 0.94, p = 0.025), LAcbShL (Cohen’s d = 1.04, p = 0.017) or LO_R_ (Cohen’s d = 1.10, p = 0.0090). Data are represented as mean ± SEM. * p < 0.05; ** p < 0.01; *** p<0.001; **** p<0.0001. A32, Prelimbic area; AcbC, accumbens nucleus, core region; CPu, caudate putamen; FrA, frontal association cortex; HP, hippocampus; LAcbSh, lateral accumbens, shell region; LO, lateral orbital cortex; MO, medial orbital cortex; mt, mamillothalamic tract; ORB, orbital cortex; Po, posterior thalamic nuclear group; PtA, parietal association cortex; S1BF, primary somatosensory cortex, barrel field; S1DZ, primary somatosensory cortex, dysgranular zone; S1Tr, primary somatosensory, trunk; S1ULP, primary somatosensory cortex, upper lip; S2, secondary somatosensory cortex; Sub, submedius thalamic nucleus; V1: primary visual cortex; V2L, secondary visual cortex, lateral; V2M, secondary visual cortex, medial; VL, ventrolateral thalamic nucleus; VM, ventromedial thalamic nucleus; VO, ventral orbital cortex; VPL, ventral posterolateral thalamic nucleus; VPM, ventral posteromedial thalamic nucleus.

Animals successfully learned the 1-Day APA task (Fig. 5). The consistent improvement over the 5 training trials in the 2 negative control groups showed that the surgery itself did not affect spatial learning, based on comparison of the first and last trials (t = 6.06, p = 0.0038 for VPM_R_; t = 8.55, p = 0.0010 for FrAR; Fig. 5c). After receiving CNO for one week, the mice in the negative control (FrA_R_ and VPM_R_) or in the CNO control group (Fig. 5d) showed intact memory recall in the probe test when compared to their own last training trial (T5) (t = 2.28, p = 0.085 for VPM_R_; t = 1.63, p = 0.18 for FrAR; t = 1.76, p = 0.12 for CNO control). In contrast, the N_shock_ was significantly increased after inhibiting each of the common hubs, S1BF_L_ (t = 4.23, p = 0.0017), V1_R_ (t = 3.76, p = 0.0045) and S2_R_ (t = 3.57, p = 0.0091), and the top or middle ranking integrator hubs, LAcbShL (t = 3.85, p = 0.0084), LO_R_ (t = 3.63, p = 0.0084) and VM_L_ (t = 2.49, p = 0.034), during the consolidation period (Fig. 5f, h). No difference was found after inhibiting a low-ranking node, the S1TrR (t = 1.14, p = 0.30). Compared to the CNO control, a significantly larger ΔN_shock_ between T5 and the probe test was found when inhibiting S1BF_L_ (Cohen’s d = 0.76, p = 0.046), V1_R_ (Cohen’s d = 0.78, p = 0.045), S2_R_ (Cohen’s d = 0.94, p = 0.025), LAcbShL (Cohen’s d = 1.04, p = 0.017) or LO_R_ (Cohen’s d = 1.10, p = 0.0090), but not VM_L_ (Cohen’s d = 0.40, p = 0.20) or S1TrR (Cohen’s d = −0.045, p = 0.47) (Fig. 5i). The T_enter_ also exhibited similar trends in these regions except for V1_R_ (Supplementary Fig. S6). These results demonstrate that inhibition of common hubs or high-ranking integrator hubs can impair memory consolidation.

## DISCUSSION

Defining brain regions and their functional involvement in the spontaneous, brain-wide reorganization that occurs after learning is essential for understanding the circuitry and mechanism of memory consolidation. Although spontaneous network activity presented in the RSNs has been identified for decades and proposed to play a role in learning and memory, this function has not been directly demonstrated. Here we find that, in addition to the HPF and mPFC, sensory areas are commonly involved following APA learning and that prefrontal, striatal and thalamic areas are pivotal for network integration. We confirm that inhibition of these RSN hubs after successful learning impairs memory formation. Our results demonstrate a causal link between post-encoding RSNs and memory consolidation, and reveal a distributed network in mediating this process, as well as providing effective methods for inferring causal hubs of behavior. This expands our understanding of the brain-wide network involved in memory formation. Considering the comparable organization and properties of human and rodent RSNs ^62,63^, our validated approaches have potential for identifying targets for intervention to enhance cognition and behavior.

The HPF and mPFC have been the most common targets in memory research. Apart from these areas, we verified the engagement of an extended network that commonly supports systems consolidation. We found that several subcortical areas in the thalamus and basal ganglia were invoked by both APA tasks, consistent with the highly distributed subcortical engrams reported in a recent study of contextual fear conditioning ^8^. Despite only a few neocortical connections being found, their hubs are required in memory consolidation. We discovered new roles of sensory areas (V1, S1BF and S2) in systems consolidation. The early visual cortex has been reported to play a role in consolidating visual working memory ^64^; however, its involvement in long-term memory consolidation has not been demonstrated. Although the hippocampus does not directly drive primary sensory areas, replay of maze-running activity patterns during slow-wave sleep has been observed in V1 ^65^, supporting its involvement. S1 has been found to engage in motor, but not spatial, memory consolidation ^66^. Recently spatially selective activity, similar to the place cells in the hippocampus, was found in S1, providing a mechanism for location-body coordination ^67^. To our knowledge, our result is the first evidence showing the involvement of S2 in memory consolidation, likely due to its role in integrating somatosensory information involved in the foot shock. Our analysis indicated that S2 is functionally connected to the CA3 region of the hippocampus, warranting further investigation of its interaction with the HPF in spatial memory formation.

Our results demonstrate the essential roles of integrator hubs in memory formation and support the notion that network integration is a key factor in memory processes. This is consistent with a rodent study which demonstrated that inhibition of brain regions estimated from covariate *c-fos* activity networks led to a reduction in the giant component correlating with the behavioral impairment ^9^. We also showed that CI is more efficient than centrality in identifying integrator hubs. In particular, there is a graded behavioral effect with both high-ranking integrator hubs (LAcbSh and LO) having large effect sizes, whereas a mid-ranking hub (VM) had a moderate effect size and a low-ranking hub (S1Tr) had minimal effect. This indicates that our analysis can predict the behavioral effects. Among the integrator hubs tested, the identification of LAcbSh is consistent with its involvement in learning and memory (for review see ^68^) and the integration of spatial information ^69^. It is also a hub that is active in both APA tasks (Fig. 2). LO is a critical prefrontal region for both decision making and the acquisition of hippocampusdependent memories ^70–72^, but its role in memory consolidation is less understood. VM, part of the motor thalamus, is the site of convergence of sensory (including nociceptive) and motor information and projects to the neocortex, particularly the mPFC ^73,74^. It is involved in decision making but its role in learning and memory remains unclear.

Systems consolidation can last for weeks, months or even years ^75,76^. How these networks transform and interact with each other over time remains unclear. Non-invasive rsfMRI allows longitudinal imaging in both animal models and humans to complement invasive imaging, such as *c-fos* imaging, which captures a snapshot during memory encoding or recall ^8,9^. The changes in RSNs and their hubs that we observed across two time points support this ongoing plasticity. Among the common hubs, the primary sensory areas were identified on post-training day 1 whereas S2 was identified on post-training day 8. This suggests a transition from primary areas during early memory consolidation to association areas later in this process. On the other hand, the integrator hubs transit from the HPF and subcortical areas on post-training day 1 to neocortical areas on post-training day 8. This is consistent with the gradual reduction in the involvement of the HPF in systems consolidation ^2,41^.

The hubs identified during the post-encoding period could be involved in forming, storing or recalling memory. We only inhibited the hubs after learning until one day before the probe test. This allowed us to determine their involvement in memory consolidation without affecting memory recall. Whether these hubs are also regions for memory storage will require further investigation. For instance, a recent study using activity tagging techniques reported that the ventrolateral orbital cortex, but not the sensory cortex, could store the contextual fear engram ^8^. Future studies may combine similar techniques to determine the specific role of RSN hubs in memory storage.

We conducted the rsfMRI in this study using a sedative protocol that is reliable in detecting RSNs ^77–79^ and post-encoding plasticity ^21,22^. Our previous study also showed that the sedative did not affect memory consolidation ^80^. Nonetheless, the detection of certain networks, such as the amygdala, which is involved in the aversive learning paradigms, may be affected ^81^. Ultrafast fMRI allows improved sensitivity for the ventral part of the brain ^82^ and enables us to detect FC with amygdala nuclei, such as the PLCo (Fig. 1). Further development of awake imaging would facilitate more comprehensive mapping and testing of functional networks not only post-encoding but also during learning or recall.

## METHODS

### Animals

121 male C57BL/6 mice (10 - 16 weeks old) were used in the experiments. Animals were housed in transparent cages and maintained on a 12 h light-dark cycle (lights on at 7a.m. and off at 7 p.m.). Food and water were provided *ad libitum.* Experiments were performed during the light phase. All experimental procedures were approved by the Animal Ethics Committee of the University of Queensland and conducted in compliance with the Queensland Animal Care and Protection Act 2001 and the Australian Code of Practice for the Care and Use of Animals for Scientific Purposes.

### Experimental design

Two sets of animal experiments were conducted, one for hub identification (Fig. 1b,c) and the other for hub verification (Fig. 1d). Four groups of mice were used for hub identification: 1-Day APA (n = 10), 1-Day control (n = 9), 5-Day APA (n = 7), and 5-Day control (n = 5). 10 groups were used for hub verification: 3 groups for the common network, 4 groups for network integration, 2 negative controls (see surgical section for details) and one CNO control (n = 9) in naïve mice without surgery.

We used DREADDs for targeted inhibition (see surgical section for detail). 4 weeks after the surgery, animals were trained in the 1-Day APA task. Immediately after they finished the last training trial (T5), they were given a water-soluble clozapine *N*-oxide (CNO dihydrochloride; cat #6329, Tocris Bioscience) via intraperitoneal (i.p.) injection (1 mg/kg dissolved in saline), followed by CNO (1 mg/kg/day) dissolved in their water to continuously suppress the network hub until one day before the probe test. This one-day interval allowed the CNO to be cleared from the body, thereby minimizing its interference in the probe test ^60^. Memory retention in the probe test was used to examine whether memory consolidation was affected by the hub inhibition.

### Behavior

In the APA task, an animal stands in a rotating circular arena (diameter: 0.9 m; rotation speed: 1 rpm) with 4 pictures as spatial cues on each side of the wall (APA equipment: Bio-Signal Group). Once the animal enters an invisible sector (shock zone) that is stable in relation to a spatial cue, a mild electric shock (0.5mA, 60Hz, 500ms) is administered. The animal needs to learn to use the visual cues to identify the exact location of the aversive zone and to avoid it. Two training protocols were used:

i. In the 1-Day APA, animals received five 10 min training sessions with an intersession interval of ~1 h completed in one day.
ii. In the 5-Day APA, animals received one 10 min training session each day for 5 consecutive days.

Training started with a habituation session (15 min) one day before the training, during which the animal did not receive any shock. Nine days after the last training day, a 10 min probe test was performed to measure memory retention in the same environment. A foot shock was delivered when the animal entered the aversive zone in the probe test. During each training or probe session, behavior was recorded by a video camera. Two control groups were included: 1-Day sham control and 5-Day sham control. In these groups, animals went through the same APA procedure as the experimental group but did not receive any foot shocks. To ensure consistency, all the behavioral experiments were started at the same time of the day. For data analysis, the number of shocks and the time to first entrance of the shock zone were analyzed by Bio-Signal Track software. Repeated measures one-way ANOVA was performed using Prism (GraphPad Software LLC).

### MRI

MRI was conducted on a 9.4 T system (BioSpec 94/30, Bruker BioSpin MRI GmbH). Two rsfMRI scan sessions were performed on each animal for hub identification. Details of the animal preparation, MRI procedure and parameter settings have been reported previously ^82^. Briefly, animals were initially anesthetized using 3% isoflurane in a 2:1 air and oxygen mixture. After being secured in an MRI-compatible holder using custom-made tooth and ear bars, a bolus of medetomidine was delivered via an i.p. catheter (0.05-0.1 mg/kg) and the isoflurane level was progressively reduced to 0.25-0.5% over 10 min, after which sedation was maintained by a constant i.p. infusion of medetomidine (0.1mg/kg/h) using a syringe pump. Key physiological parameters, including arterial oxygenation saturation (SpO2), rectal temperature, heart rate and respiratory rate, were measured by an MRI-compatible monitoring system (SAII Inc). Body temperature was maintained at 36.5°C with a heated waterbath.

After high-order shimming, structural T2-weighted MRI (resolution = 0.1 × 0.1 × 0.3 mm^3^) and the visual task were first conducted to ensure optimal physiology and neurovascular coupling. A flashing blue light at 5Hz was delivered by an optical fiber in a block design with 21s on and 39s off. The rsfMRI scan was then acquired using multiband gradient-echo echo-planar imaging ^82^ with TR/TE = 300/15 ms, thickness = 0.5 mm, gap = 0.1 mm, 16 axial slices covering the whole cerebrum with in-plane resolution of 0.3 × 0.3 mm^2^. 2000 volumes were acquired in 10 min and repeated 3 times with an inter-run interval of 2 min. To ensure consistency, the rsfMRI scan started ~45min after the bolus injection of medetomidine.

### Surgical procedure for DREADDs

Surgeries were performed at least one month before behavioral training. The animal was anesthetized with 1.5-2% isoflurane during surgery. Enrofloxacin (6 mg/kg) and carprofen (5 mg/kg) were injected subcutaneously to prevent infection and relieve pain and inflammation, respectively. Body temperature was maintained at 37 °C with a heating pad. During surgery, 0.25-0.3 μL of virus (AAV2/1-pSyn-hM4D(Gi)-T2A-mScarlet) was injected into the following target areas based on the coordinates (relative to Bregma) in the Paxions and Franklin Mouse Brain atlas, fifth edition.

#### Common hubs

(i) V1_R_ (n = 10). ML: −2.30mm; AP: −4.15mm; DV: −0.50mm.
(ii) S2_R_ (n = 8). ML: −3.70mm; AP: −0.71mm; DV: −1.40mm.
(iii) S1BF_L_ (n = 11). ML: +2.88mm; AP: −1.07mm; DV: −0.85mm.

#### Integrator hubs

(iv) LAcbSh_L_ (n = 7). ML: +1.70mm; AP: +1.10mm; DV: −3.85mm.
(v) LO_R_ (n = 8). ML: −1.65mm; AP: +2.57mm; DV: −1.9mm.
(vi) VM_L_ (n = 10). ML: +0.75mm; AP: −1.18mm; DV: −4.05mm.
(vii) S1Tr_R_ (n = 7). ML: −1.63mm; AP: −1.60mm; DV: −0.57mm.

#### Negative control

(viii) VPM_R_ (n = 5). ML: −1.5mm; AP: −1.80mm; DV: −3.3mm.
(ix) FrA_R_ (n = 5). ML: −1.70mm; AP: +3.00mm; DV: −0.50mm.

Virus was injected using a Nanoject III (Drummond Scientific) with a slow injection rate (0.03 μL/min) over 10 min. The glass pipette was retained in place for another 6 min and then slowly retracted. After injection, the wound was closed using Vetbond (3M) and sutured. Enrofloxacin and carprofen were administered for another two days. Animals were kept in their home cage (group housing of 2-4 animals per cage) for 4 weeks to recover and to allow expression of the virus before APA testing.

### Behavior and CNO treatment

The same 1-Day APA task was used for spatial memory training. Immediately after the 1-Day APA training, water-soluble CNO was administered to the animals (1mg/kg, i.p.) to inhibit the neuroactivity of the target brain regions. Water containing CNO (1mg/kg/day) was then provided for 7 days to keep the target brain areas inhibited during memory consolidation. This was replaced with normal water 24 h before the probe test to minimize the effects of CNO on behavioral performance. On post-training day 8, a 10 min probe trial was performed to test memory retention.

### Histology

Mice were administed an overdose of sodium pentobarbitone and transcardially perfused with 40 ml of phosphate-buffered saline (PBS), followed by 45 ml of 4% paraformaldehyde in PBS for fixation. The brain was extracted and fixed at 4°C for 12-24 h. It was then washed once with PBS and transferred to a 30% sucrose solution for 36 h prior to sectioning. 40 μm thick sections were cut using a sliding microtome and collected in a 1:6 series. Cell nuclei were stained by 4’,6-diamidino-2-phenylindole (DAPI, catalog #6329; Sigma Aldrich). Sections were first washed once in PBS for 10 min, and then incubated in 1:5000 DAPI-PBS solution for 15 min at room temperature. After two washes, the sections were mounted on SuperFrost slides using fluorescence mounting medium (Dako, Agilent). Images were captured using a slide scanner (Metafer VSlide Scanner, MetaSystems) and microscope (Axio Imager Z2, Zeiss) with a 20x 0.8 NA / 0.55mm objective lens.

### rsfMRI data processing

The rsfMRI data were processed using MATLAB (MathWorks Inc), FSL (v5.0.11, https://www.fmrib.ox.ac.uk/fsl), AFNI (ver 17.2.05, National Institutes of Health, USA) and ANTs (v2.3.1, http://stnava.github.io/ANTs). The k-space data of the multiband EPI were first phase-corrected and reconstructed in MATLAB. After motion correction by FSL mcflirt, the geometric distortion was corrected by FSL TOPUP. The brain mask was extracted automatically using PCNN3D ^83^, followed by manual editing. Nuisance signals, including quadratic drift, 6 motion parameters and their derivatives, 10 principal components from tissues outside the brain, and mean signal of the cerebrospinal fluid, were then regressed out ^84^. The data were band-pass filtered at 0.01-0.3 Hz to account for any potential frequency shift under sedation. This frequency range could also remove the aliased respiratory and cardiac signal variations in the high sampling rate data. The rsfMRI was coregistered to an EPI template by linear and nonlinear transformations using ANTs. The data were then smoothed by a 0.6 mm Gaussian kernel.

Seed-based correlation analysis was used to measure FC across the brain. Based on the Australian Mouse Brain Mapping Consortium (AMBMC) atlas (https://imaging.org.au/AMBMC/AMBMC), the brain was divided into 230 bilateral regions of interest (ROIs). The DSURQE atlas (https://wiki.mouseimaging.ca/) was used to label regions not yet defined in the AMBMC atlas, such as the amygdala, hypothalamus, midbrain and brainstem (pons). The mean time-series of each brain region was extracted as a seed signal. Pearson’s correlation coefficients between seed time-courses were calculated using AFNI 3dNetCorr. Fisher’s *z*-transformation was used to convert correlation coefficients to *z* values. Connectivity matrices from the 3 repeated scans were calculated for each animal. Quality control (QC) was conducted based on the presence of visual task activation. If the visual activation was not detectable, the physiological condition and neurovascular coupling were regarded as sub-optimal, and the scan was discarded. Based on this criterion, 24% of scans were discarded (Supplementary Table S3). The matrices of each animal at each time point that passed the QC were averaged.

Between-group differences were calculated by two-sample t-test and thresholded at p<0.05 (FDR corrected) using the Network Based Statistics toolbox (https://sites.google.com/site/bctnet/comparison/nbs). For hub identification, a lower threshold (uncorrected) was used to generate a weighted (t-score) network matrix.

### Graph theory analysis

To characterize the RSNs, the following graph theory parameters were calculated by the Brain Connectivity Toolbox (http://www.brain-connectivity-toolbox.net) and the graph functions in Matlab:

Global efficacy:

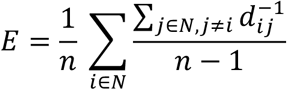

where *d_ij_* is the shortest path length between nodes *i* and *j*, and *N* is the total number of nodes. A value of 1 indicates maximum efficiency.

Modularity was calculated using the Newman’s spectral community detection algorithm:

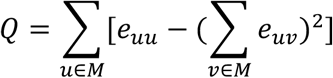

where the network is fully subdivided into a set of nonoverlapping modules *M*, and *e_uv_* is the proportion of all links that connect nodes in module *u* with nodes in module *v.* The higher the Q value, the larger degree of network segregation.

Degree centrality of a node *i*:

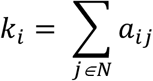

where *a_ij_* is the connection between nodes *i* and *j*. *a_ij_* = 1 when a link (*i, j*) exists, and *a_ij_* = 0 otherwise (*a_ii_* = 0 for all *i*).

Closeness centrality of node *i*:

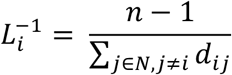

Betweenness centrality of node *i*:

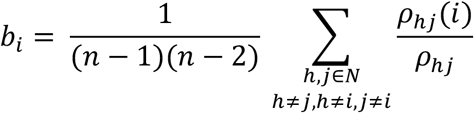

where *ρ_hj_* is the number of shortest paths between nodes *h* and *j*, and *ρ_hj_*(*i*) is the number of shortest paths between h and *j* that pass through *i*.

Eigenvector centrality of node *i*:

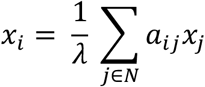

where *λ* is a constant and *x* is the eigenvector of the binarized network matrix.

The giant component is defined as the largest connected component in a network ^9^. It can be represented by the number of nodes in the largest connected component. The ratios of the giant components in the CI analysis were used to represent the change in the giant component when nodes were removed from the network.

### FC – behavior correlation

We correlated the strength of each significant FC with two behavioral indices (the number of shocks and the time to first entrance of the shock zone) in the probe test using Pearson’s correlation coefficient, with p < 0.05 (two-tailed) regarded as significant. After correlation analysis, we sorted the significant connections based on their absolute value of the correlation coefficient.

### Common networks between 1-Day and 5-Day APA

We detected overlapping connections from the network matrix obtained from the two-sample t-test between the APA and control groups (p < 0.05, uncorrected). From the overlapping connections, we calculated FC–behavior correlations and sorted (ranked) the significant connections (p < 0.05) by their absolute value of behavior correlation.

### CI analysis

CI analysis was applied to the between-group difference network using an optimized implementation ^59^ (https://github.com/zhfkt/ComplexCi/releases) that calculates the value of each node with the following formula:

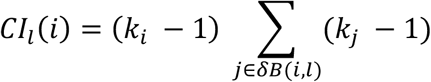

where *k_i_* is the degree of node *i*, and *δB*(*i, l*) is the frontier of the ball of radius *l* which is the set of nodes at a particular distance from *i*. The value is calculated iteratively by removing nodes until all nodes in the network are eliminated ^57^. In this analysis, we used the ball radius *l* = 2 as we found that a larger radius gave nearly the same results. We ranked the nodes according to how fast the size (number of nodes) of the “giant component” collapsed by removing the selected node. To find the reliable ranking list, we calculated the mean CI ranking of each node under three thresholds (p < 0.05, p < 0.01, p < 0.005) and sorted the node by the mean CI rank value.

### Null distribution of network property analysis

To determine whether the difference matrices between the APA and control groups were significantly different from random networks, 5000 random networks were created using the function “null_model_und_sign” of the Brain Connectivity Toolbox that preserves the degree and strength distributions of the real network. For each of the 5000 random networks, curves of three network properties, including the giant component, global efficiency and modularity were generated under thresholds ranging from t = 2 to 3.8 (step of 0.2). The AUC of each curve was calculated to form the null distribution for each network property.

### Null distribution of common network analysis

To estimate the null distribution of the common network, 5000 permutations of FC matrices were created by randomly assigning each individual FC matrix into the APA groups and controls. Between-group differences of these permutated matrices were tested by two-sample t-tests and thresholded at p < 0.05. The common FC of the permutated 1-Day APA and 5-Day APA data was correlated with the N_shock_ or T_enter_ of the probe test and thresholded at p < 0.05. The number of connections surviving these thresholds from the 5000 permutations formed the null distribution of the common network detection (Supplementary Fig. S3a).

## Acknowledgement

This study was funded by Australian Research Council Discovery Project grant #180103319. Z.L. was supported by a Research Training Program scholarship of the University of Queensland. We thank Ms Rowan Tweedale for proof reading and Prof Ethan Scott for helpful discussion.

## Author contribution

K.C., P.O. and P.S. contributed to designing experiments.

Z.L. H.L. and D.A. conducted experiments.

Z.L. and K.C. conducted data analysis.

Z.L., P.O., P.S. and K.C. wrote the manuscript.

